# Alternative polyadenylation mediated by CPSF5 regulates pro-inflammatory gene expression

**DOI:** 10.64898/2026.01.31.703030

**Authors:** Matthias Lemmer, Doris Lindner, Georg Stoecklin, Johanna Schott

**Affiliations:** Mannheim Institute for Innate Immunoscience (MI3) and Mannheim Cancer Center (MCC), Medical Faculty Mannheim, Heidelberg University, Mannheim, Germany; Center for Molecular Biology of Heidelberg University (ZMBH), German Cancer Research Center (DKFZ)-ZMBH Alliance, Heidelberg, Germany

**Keywords:** alternative polyadenylation, CFIm25, CPSF5, macrophages, inflammation

## Abstract

The majority of mammalian genes contains more than one functional cleavage and polyadenylation site, so that selection of alternative sites leads to different 3’ ends of the mature transcripts. This mechanism can lead to the exclusion or retention of regulatory sequence elements, which affects post-transcriptional regulation of gene expression, such as RNA stability, localization or translation efficiency. We used 3’ end sequencing to assess alternative polyadenylation during the response of macrophages to LPS, and observed a strong global shift towards proximal polyadenylation sites. This was accompanied by a decreased expression of cleavage and polyadenylation factors, including CPSF5, which is known to favor selection of distal polyadenylation sites. Upon depletion of CPSF5 in macrophage, we observed global transcript shortening, and an induction of TNF and other pro-inflammatory cytokines without LPS-stimulation. Analysis of RNA-seq data from monocytes of sepsis patients revealed that CPSF5 expression and alternative polyadenylation are also affected *in vivo*. Usage of distal polyadenylation sites showed a negative correlation with *TNF* mRNA expression in human monocytes. Our data suggest that transcript shortening mediated by CPSF5 repression contributes to the induction of pro-inflammatory genes.

## Introduction

Gene expression in macrophages is highly dynamic and extensively controlled at the post-transcriptional level, which allows a rapid but temporally limited response to inflammatory stimuli (reviewed in [1]). Many regulatory sequence motifs that control mRNA stability and translation efficiency are located in the 3’ UTR, which serves as a platform for binding of miRNAs or RNA-binding proteins. A set of structured hairpin elements, for example, plays a prominent role in the immune system by repressing stability or translation of mRNAs that encode for inflammatory factors [2, 3]. AU-rich elements (AREs) are also enriched in immune-relevant genes and can recruit a variety of RNA-binding proteins with different effects on mRNA stability or translation (reviewed in [4]). In addition, several miRNAs have been shown to regulate immune cell function, for example by affecting the expression of components of the TLR4 signaling cascade (reviewed in [5]).

The 3’ end of transcripts is determined by cleavage of the immature transcript about 15 -30 nt downstream of the polyadenylation signal, which is typically an AAUAAA hexamer or variants of this motif. The polyadenylation signal is recognized by the cleavage and polyadenylation specificity factor (CPSF) complex. The cleavage stimulation factor (CstF) and the mammalian cleavage factor II (CFIIm) bind to elements downstream of the polyadenylation signal and lead to the activation of the endonuclease CPSF73, which cleaves the pre-mRNA at the polyadenylation site (PAS). The poly(A) polymerase (PAP) then adds the poly(A)-tail (reviewed in [6]).

Many genes, however, have more than one polyadenylation signal. Selection of a PAS can be regulated by the CFIm complex, which recognizes UGUA motifs upstream of the polyadenylation signal, favors the selection of distal PASs and thereby induces transcript lengthening. In contrast, the CFIIm component PCF11 increases the usage of proximal PASs, leading to shorter transcript isoforms [7]. Many other factors have been described to affect PAS selection. These include RNA binding proteins (RBPs), RNA modifications, post-translational modifications of the cleavage and polyadenylation machinery, transcription factors, chromatin conformation, and DNA modifications (reviewed in [6]). These do not seem to systematically favor the selection of proximal or distal PASs, but rather affect PAS usage in a gene-specific manner [7].

By alternative polyadenylation (APA) of protein-coding genes, different 3’ UTR isoforms are generated. Derti et al. reported that about 70% of all protein-coding human genes undergo tissue-specific APA [8], while Hoque et al. identified multiple PASs in 79% of protein-coding mouse genes [9]. APA can lead to the retention or exclusion of regulatory elements and thereby affect post-transcriptional control. Activation of CD4^+^ T-cells, for example, induces the loss of miRNA binding sites on many mRNAs by global 3’ UTR shortening, which enhances protein expression [10].

Regulation of PAS selection, in a global or gene-specific way, has been described in many different conditions, including development, cancer and immunity. In this study, we addressed the role of APA in the reaction of macrophages to LPS.

## Materials and methods

### Cell culture

RAW264.7 cells were cultured at 37°C in a humidified atmosphere with 5% CO_2_ in Dulbecco’s modified Eagle’s medium (DMEM, Gibco) supplemented with 10% (v/v) fetal bovine serum (FBS, Biochrome), 2 mM L-Glutamine, 100 U/ml penicillin and 0.1 mg/ml streptomycin (all PAN Biotech). For splitting, cells were detached every two to three days by scraping in warm PBS and diluted 1:10.

### QuantSeq 3’ mRNA-sequencing

RAW264.7 cells were treated with 100 ng/ml LPS (E. coli O111:B4, Sigma). Total RNA was isolated using the Universal RNA Purification Kit (Roboklon) including an on-column DNA digest step according to the manufacturer’s instructions. Sequencing libraries were prepared with the QuantSeq 3’ mRNA-Seq V2 Library Prep Kit REV (Lexogen) following the manufacturer’s instructions with 500 ng of total RNA. Libraries were sequenced as equimolar pools on a Nextseq550 at the Next Generation Sequencing core facility of the Medical Faculty Mannheim.

Read sequences were aligned with bowtie2 to the mouse genome (mm39) [11]. Read coverage was calculated with the genomecov function of bedtools [12], and PASs were identified using an in-house developed R script. PASs were annotated to genes using the basic set of transcripts from Gencode VM32 as downloaded from the UCSC Genome Browser (wgEncodeGencodeBasicVM32). Only PASs were considered that had a sum of read counts ≥ 120 across all samples, and a relative usage of at least 20% in one of the samples. Changes in PAS usage were analysed using the R package DEXseq [13]. DESeq2 [14] was used to analyse differential gene expression after summing up the read counts of all PASs per gene.

### Western blotting

For measuring the expression of CPSF proteins after treatment with LPS, cells were washed twice with ice-cold PBS and lysed in RIPA buffer (50 mM Tris pH 8.0, 150 mM NaCl, 1% NP-40, 0.5% sodium deoxycholate, 0.1% SDS) supplemented with cOmplete EDTA-free Protease Inhibitor Cocktail (Roche). Lysates were incubated for 30 min on ice, and cell debris was removed by centrifugation for 5 min at 13.000 × g at 4°C. Samples were denatured for 10 min at 95°C in SDS sample buffer (SB; 50 mM HEPES pH 7.4, 2% SDS, 10% glycerol, bromphenol blue, 50 mM dithiothreitol). For measuring CPSF expression upon shRNA-mediated knockdown, cells were directly lysed in 2 × SB buffer and sonicated using the Bioruptor system according to the manufacturer’s instructions. Proteins were then separated on a 5-20% polyacrylamide gradient gel. Electrophoresis was performed in 1 × running buffer (25 mM Tris, 250 mM glycine, 0.1% SDS) with 30 mA per gel. Proteins were then transferred onto a 0.2 μm pore size nitrocellulose membrane in 1 × blotting buffer (20 mM Tris, 150 mM glycine, 20% EtOH) at 100 V and 4°C for 90 min using the wet blotting technique. Protein loading and blotting efficiency were assessed by Ponceau S staining of the membrane. Destaining was performed with TBS-T (50 mM Tris-HCl pH 7.5, 150 mM, NaCl, 0.1% Tween-20), and blocking with 3% BSA in TBS-T for 1 h at RT. Membranes were incubated with primary antibodies (1:1,000; ACTB: Abcam ab8227-50, CPSF5: Santa Cruz sc-81109, CPSF6: Abcam ab99347) by shaking overnight in 1 × PBS/0.02% sodium azide. Membranes were washed at least three times for around 15 min with TBS-T before incubating with α-rabbit horseradish peroxidase-coupled secondary antibodies (Jackson ImmunoResearch) diluted 1:5,000 in 1 × TBS-T for 1 h at RT. Membranes were washed at least three times for around 15 min with TBS-T before protein signals were detected using Clarity ECL Western Blotting Substrates (Bio-Rad) according to the manufacturer’s instructions and a ChemiDoc System.

### Lentiviral transduction with shRNAs

Plasmids expressing shRNAs were purchased as commercially available bacterial glycerol stocks from Sigma-Aldrich (anti-*Cpsf5*: TRCN0000109258). Lentiviral particles were produced by seeding 3 × 10^6^ HEK-293T cells in 8 ml regular medium and incubated overnight. One hour before transfection, the medium was exchanged with 6 ml of fresh medium. The transfection mix was prepared in 1.5 ml Opti-MEM (Thermo Scientific) by mixing 6.42 µg of plasmid pCMV Δ8.91, 2.16 µg of plasmid pMD.G and 6.42 µg of the shRNA-expressing plasmid with 45 µl TurboFectin 8.0 (OriGene). The reaction mix was thoroughly vortexed and incubated for 20 min at RT. Afterwards, the mix was added dropwise to the HEK-293T cells. After 48 h, the supernatant was harvested, filtered through a 0.45 filter. and immediately stored at -80°C. The virus-containing supernatant was diluted 1:2 in regular medium. After 48 h, the medium was removed and cells were re-seeded for the following experiment.

### Measurement of cytokine production

Cells were seeded after 48 h of control or *Cpsf5* knockdown (see above) and treated with 100 ng/ml LPS. Supernatants of stimulated macrophages were collected and stored at -80°C. TNF protein concentrations were measured using the ELISA Pro: Mouse Tnf-α Kit (Mabtech) according to the manufacturer’s instructions. For cytokine arrays, the Proteome Profiler Human Cytokine Array Kit (R&D) was used according to the manufacturer’s instructions.

### Analysis of distal 3’ UTR usage in RNA-seq data

Fastq files of CD14^+^ monocytes of the conditions healthy, critically ill, early and late sepsis generated by Washburn et al. [15] were retrieved from ENA. The R2 reads were aligned to the human genome version hg38 using STAR [16], providing the basic transcript set of Gencode V38 as a gtf. Read count density on the most distal 200 nt of each gene was compared to the average read count density on the entire gene as a measure for distal PAS usage. For read annotation, the Bioconductor package GenomicRanges [17] and the featureCounts function of the subread package [18] were used.

## Results

### LPS-treatment leads to a global shift to proximal polyadenylation sites in macrophages

To measure PAS usage in macrophages during the response to LPS, we performed 3’ end sequencing in untreated RAW264.7 cells and after stimulation with LPS for 1 h and 16 h. Using the QuantSeq REV kit for library preparation, PASs can be precisely mapped, because the first nucleotide of the sequencing read directly corresponds to the cleavage site (Fig. 1A). Across all conditions, we detected PASs on 11,813 genes. For 34.5% of these genes, we identified more than one PAS, with two PASs on 3056 genes, three PASs on 885 genes, and more than three PASs on 137 genes (Fig. 1B). As expected, the largest proportion of these PASs (74%) is located in exonic regions of the 3’ UTR (Fig. 1C). Interestingly, many sites were identified in intronic (13.5%) or exonic (1.5%) regions of the open reading frame (ORF), which leads to the loss of the annotated termination codon and possibly to the inclusion of intronic premature termination codons. To detect specifically 3’ UTR lengthening or shortening, we focused our analysis on exonic 3’ UTR sites. After 1 h of LPS treatment, only 26 genes showed significant changes in PAS usage (Fig. 1D). On *Maff* mRNA, which is the most prominent case, usage of the distal PAS is strongly induced (Fig. 1D-E), which leads to 3’ UTR lengthening. After 1 h of LPS stimulation, there is no systematic bias for increased usage of proximal or distal PASs (shown in red and blue, respectively). After 16 h, however, there is a global increase in proximal PAS usage (Fig. 1D). Among the 1244 genes with significant changes in PAS usage, 874 displayed a significant increase for the most proximal PAS. On *Klf11* mRNA, for example, the distal PAS is used more often than the proximal PAS under control conditions, but upon 16 h of LPS treatment, almost exclusively the proximal site is selected (Fig. 1D-E).

**Figure 1.**
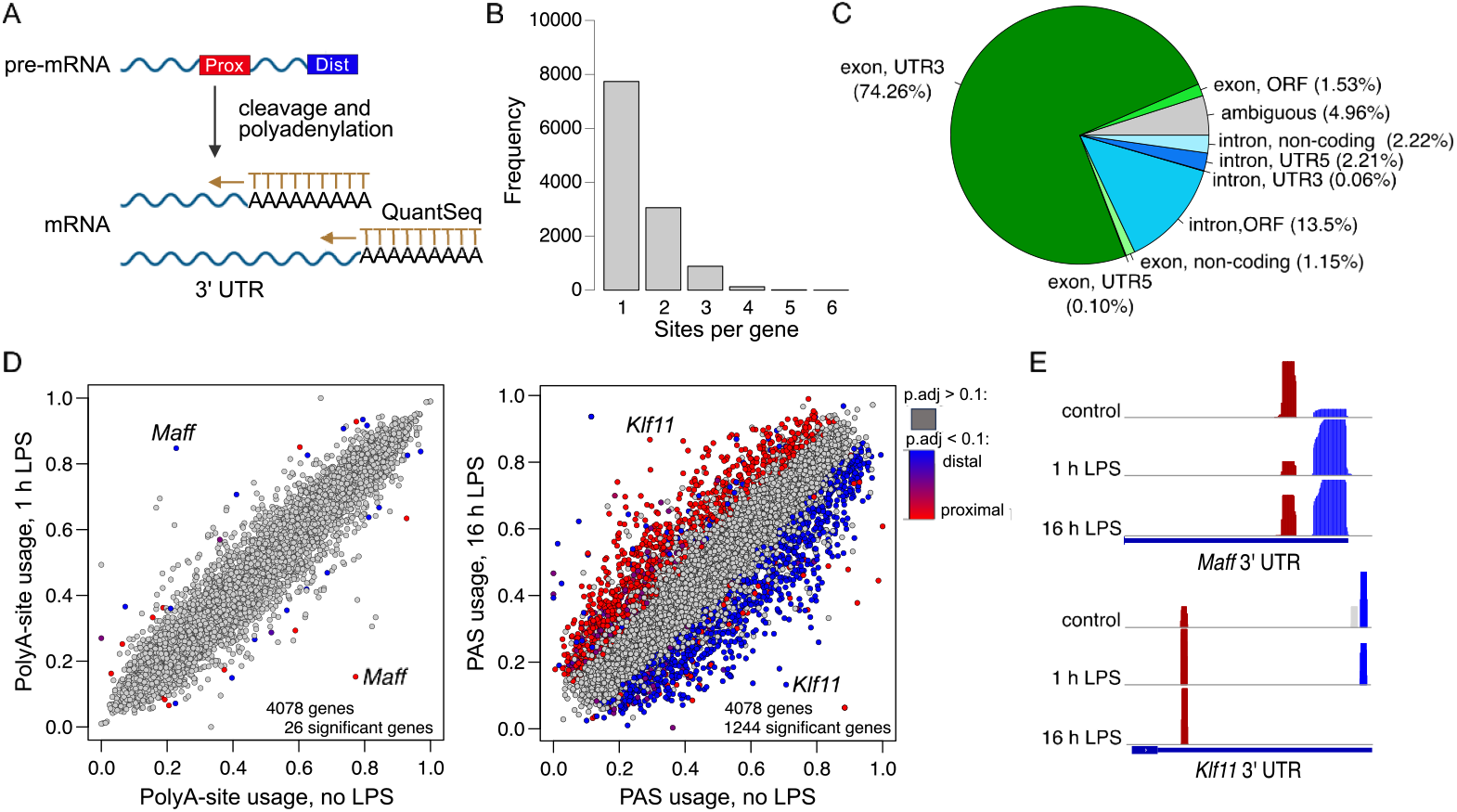
LPS-treatment induces global 3’ UTR shortening in macrophages. **(A)** QuantSeq 3’ mRNA sequencing allows a precise identification of PASs. **(B)** Number of PASs per gene as identified by QuantSeq in RAW264.7 cells. **(C)** Location of PASs according to genomic features. **(D)** Scatter plot of PAS usage for genes with at least two PASs detected by QuantSeq in RAW264.7 macrophages after 1 h or 16 h LPS treatment (100 ng/ml, n = 3) compared to control cells. PASs were filtered based on the sum of reads per site across all samples (≥ 120) and a minimum of 20% usage in at least one of the samples. Statistically significant (p.adj < 0.1) PASs are colored red (proximal PAS), blue (distal PAS) or purple (PASs between the most proximal and the most distal PAS on genes with more than two PASs). **(E)** QuantSeq read coverage on *Maff* and *Klf11* in control cells and after 1 h or 16 h of LPS treatment.

### Expression of CPSF5 is repressed by LPS-treatment

A global one-directional shift in PAS usage may be caused by reduced expression of cleavage and polyadenylation factors. Therefore, we analyzed differential RNA expression in our QuantSeq ata by aggregating the read counts of all PASs per gene. Among the detected components of the cleavage and polyadenylation machinery, *Pcf11* (encoding for a subunit of the CFIIm complex) is most strongly affected after 1 h of LPS treatment. PCF11 mostly favors the usage of proximal PASs [7]. Induction of *Pcf11* mRNA, however, was not detectable anymore after 16 h of LPS treatment, where we observed a reduced expression of several cleavage and polyadenylation factors. Among these factors, especially CPSF5 was described to drive the usage of distal PASs [7]. Together with CPSF6 and CPSF7, CPSF5 dimers form the CFIm complex and mediate binding to UGUA sequences in proximity of polyadenylation signals [6]. In contrast to *Cpsf5*, expression of *Cpsf6* and *Cpsf7* mRNAs is not reduced by LPS treatment (Fig. 2B). At the protein level, however, CPSF6 expression is also affected after long-term exposure to LPS (Fig. 2C). This may be explained by destabilization of the CPSF6 protein when the CFIm complex cannot assemble due to decreased CPSF5 abundance.

**Figure 2.**
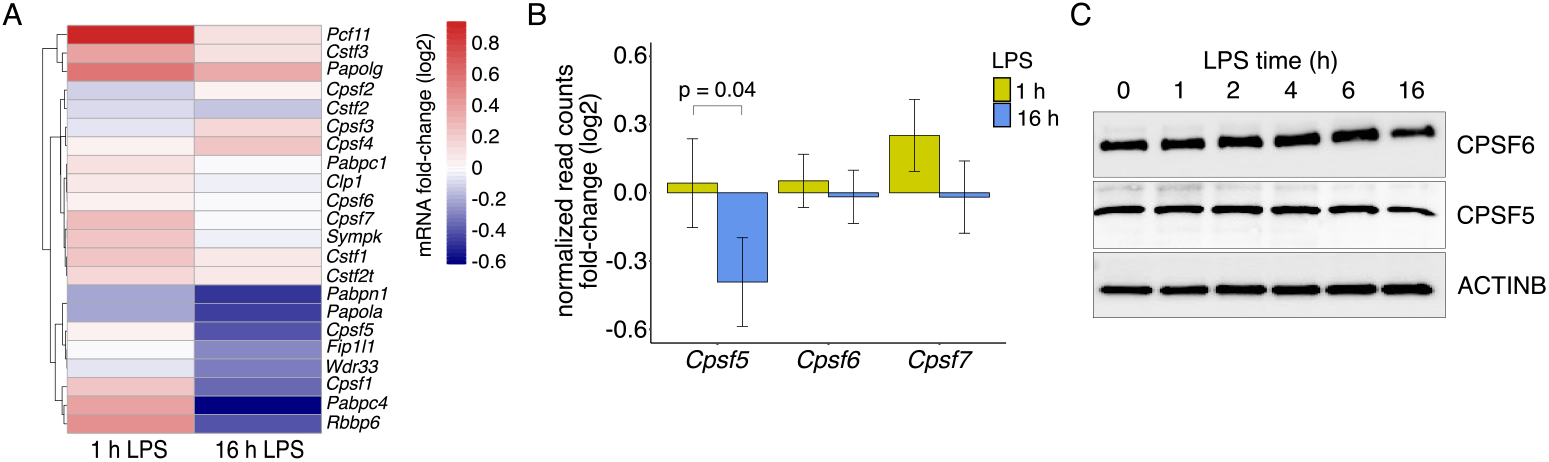
Expression of *Cpsf5* is repressed upon LPS-treatment in macrophages. **(A)** QuantSeq reads were aggregated per gene to determine differential mRNA expression. Log2-transformed fold-changes upon 1 h or 16 h LPS treatment in RAW264.7 macrophages are shown as a heatmap for components of the cleavage and polyadenylation machinery according to Ogorodnikov et al. **(B)** Expression of the three components of the CFIm complex (*Cpsf5, Cpsf6* and *Cpsf7*) at the mRNA level (log2-transformed fold-change ± standard error as determined with DESeq2). **(C)** Expression of CPSF5 and CPSF6 proteins as determined by Western blotting at the indicated time-points of LPS treatment.

### Shortening of 3’ UTRs leads to de-regulated cytokine expression

To investigate the effect of transcript shortening induced by repression of CPSF5, we performed shRNA-mediated depletion of CPSF5 in RAW264.7 cells (Fig. 3A). Mapping of PASs with QuantSeq 3’ mRNA-Seq confirmed that loss of CPSF5 leads to preferential usage of proximal PASs. In total, 2825 genes showed a significant change in PAS usage (p.adj < 0.1), and most shifts occurred from distal to proximal PASs (Fig. 3B). There was a strong overlap between APA events detected upon LPS and upon *Cpsf5* knockdown, with 77% of the LPS-regulated genes also being affected by CPSF5 knockdown (Fig. 3C). Among these genes, we identified one subunit of the nuclear factor-kappa-B (NF-κB) family of transcription factors, *Nfkb1*, which encodes for p105/p50 (Fig. 3D). *Nfkb1* shows a very similar shift from the distal to the proximal PAS, confirming that *Cpsf5* knockdown mimics the effect of LPS-induced APA. As a measure for macrophage activation, we assessed TNF secretion in the supernatants of stimulated and unstimulated macrophages using enzyme-linked immunosorbent assay (ELISA). Knockdown of *Cpsf5* alone led to a ∼5-fold induction of TNF secretion (Fig. 3E). Upon LPS-treatment, TNF secretion was similar between knockdown and control cells. To measure a broader spectrum of cytokines, we analyzed supernatants of control and knockdown cells on a cytokine array. In addition to TNF, CCL5 and CXCL2 production was strongly increased by *Cpsf5* knockdown alone, but not upon LPS treatment (Fig. 3F-G). IL6 and CSF2 were only detectable in the supernatants of LPS-stimulated macrophages, and their induction was impaired by CPSF5 knockdown (Fig. 3F-G).

**Figure 3.**
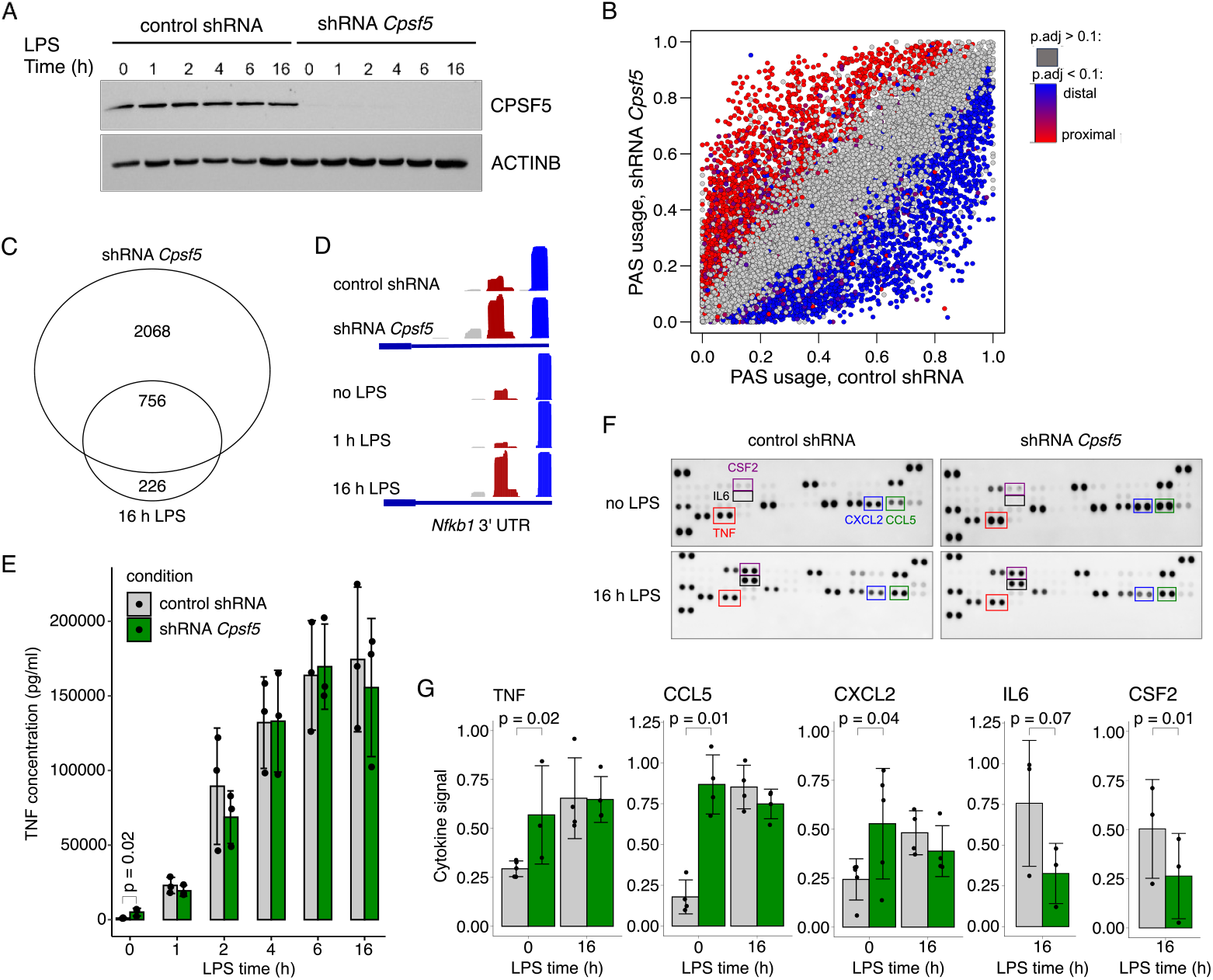
Shortening of 3’ UTRs by knockdown of *Cpsf5* leads to de-regulated cytokine expression. **(A)** Knockdown of *Cpsf5* mediated by lentiviral transduction of RAW264.7 cells with shRNA-expressing vectors. Protein levels are measured by Western blotting from samples analyzed in (E). **(B)** Scatter plot of PAS usage for genes with at least two PASs detected by QuantSeq in RAW264.7 macrophages upon transduction with control shRNA or shRNA targeting *Cpsf5* (analyzed as in Fig. 1D). **(C)** Overlap of PASs regulated upon LPS treatment and CPSF5 depletion. **(D)** QuantSeq read coverage on *Nfkb1* in control cells, after 1 h or 16 h of LPS treatment, and upon *Cpsf5* knockdown. **(E)** TNF protein levels measured by ELISA in the supernatants of RAW264.7 at the indicated time-points of LPS-treatment. **(F)** Cytokine arrays to measure protein concentrations in the supernatant of control cells and cells treated for 16 h with LPS combined with control shRNA or *Cpsf5* shRNA transduction. **(G)** Quantification of cytokine arrays as in (F). Bar plots show means ± SD. P-values were obtained from paired t-tests.

### Distal PAS usage is altered in human sepsis

Our data from RAW246.7 cells stimulated *in vitro* with LPS indicate that the inflammatory reaction of macrophages is accompanied by a repression of *Cpsf5* expression, which leads to the preferential usage of proximal PASs. To transfer these findings into an inflammatory condition *in vivo*, we reanalyzed publicly available RNA-seq data obtained from CD14^+^ monocytes during early and late sepsis, of critically ill patients and of healthy donors [15]. Expression of *CPSF5* mRNA was suppressed in all conditions compared to the healthy controls (Fig. 4A). As a proxy for distal PAS usage, we quantified the relative read density on the last 200 nt per gene, and determined the median of this value for all genes per patient (Fig. 4B). Interestingly, distal PAS usage shows a broader distribution in critically ill patients and during early sepsis than in healthy donors. This indicates that PAS usage is altered in monocytes during inflammatory conditions *in vivo*, with a trend towards reduced distal PAS usage (Fig. 4B). Despite the reduced *CPSF5* expression, the effect on distal PAS usage does not persist during late sepsis, where the distribution of distal PAS usage is very similar to the healthy controls. Interestingly, *TNF* mRNA expression does not systematically differ between the groups (Fig. 4C). Nevertheless, *TNF* mRNA abundance correlates negatively with distal PAS usage (Pearson’s correlation coefficient: -0.3; p-value = 0.01).

**Figure 4.**
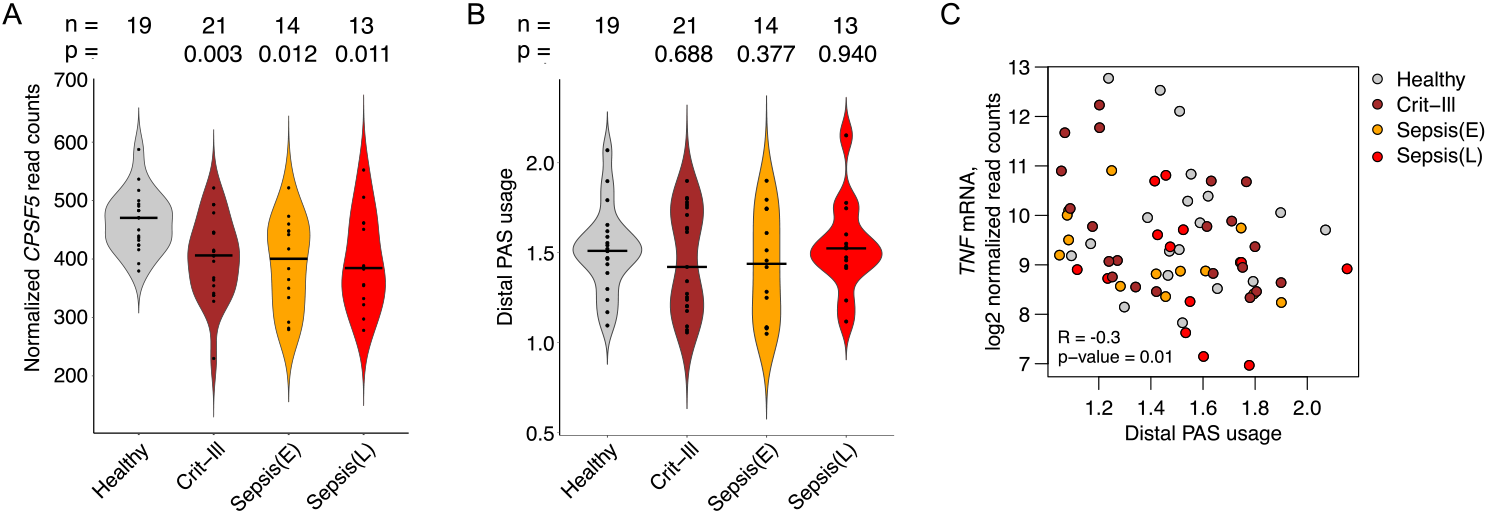
Expression of *Cpsf5* and PAS selection is affected in human sepsis. **(A)** *Cpsf5* expression in RNASeq data [15] of CD14^+^ monocytes isolated from healthy donors, critically ill patients and during early or late sepsis. P-values were determined using Wilcoxon-rank-sum tests comparing each group to the healthy controls. **(B)** Usage of distal PASs estimated from the relative read density on the last 200 nt per gene. For each patient, the median of all detected genes is shown. P-values were determined using Wilcoxon-rank-sum tests comparing each group to the healthy controls. **(C)** Correlation of *TNF* mRNA expression estimated from normalized read counts with distal PAS usage determined as in (B). R: Pearson’s correlation coefficient.

## Discussion

Using 3’ end sequencing in LPS-stimulated macrophages, we observed a strong global shift towards proximal PASs during the inflammatory response. Among the factors that regulate APA with a directional bias, *Pcf11* expression is transiently induced after 1 h, while *Cpsf5* is repressed after 16 h of LPS treatment (Fig. 2A). Global changes in PAS selection have also been described for macrophages upon infection with different pathogens. Infection of macrophages with the vesicular stomatitis virus (VSV) leads to reduced expression of cleavage and polyadenylation factors including CPSF5, and to the preferential usage of proximal PASs [19]. While Listeria and Salmonella infection likewise induce transcript shortening [20], infection with Mycobacterium tuberculosis has the opposite effect [21].

Most studies link 3’ UTR shortening to post-transcriptional induction of gene expression by increased mRNA stability or translation efficiency. Pai et al. described wide-spread evasion from miRNA-mediated repression due to the loss of miRNA target sites [20]. Jia et al. used reporter gene assays to confirm that the shorter 3’ UTR isoforms of several APA affected genes including *NFKB1* confer induction at the post-transcriptional level [19]. Likewise, Ge et al. could demonstrate enhanced mRNA stability and translation efficiency of selected genes due to 3’ UTR shortening [22]. On the other hand, isoform-specific measurements of mRNA stability and translation in mouse fibroblasts revealed only a small tendency of longer half-lives for mRNAs with shorter 3’ UTR isoforms, and greater translational efficiency [23]. The impact of 3’ UTR isoform choice may depend on the cell line or tissue, as different mechanisms of gene expression control may be more or less prominent in different cellular contexts. In addition, some regulatory motifs also enhance mRNA stability or translation efficiency, as described for the ARE-binding protein HuR (reviewed in [4]). Finally, 3’ UTR shortening does not only lead to the loss of sequence elements, but might also activate elements that are otherwise located in the center of the 3’ UTR [24]. For both miRNA target sites and AU-rich elements, the position within the 3’ UTR has been described to affect their function and activity [25, 26]. Beyond the impact on post-transcriptional regulation mediated by 3’ UTR elements, APA can affect gene expression by loss of the canonical stop codon. We observed that 13.5% of the PASs in macrophages are located in intronic regions upstream of the stop codon (Fig. 1C). These events can lead to the synthesis of protein isoforms that are truncated at the C-terminus [27], because the ribosome is very likely to encounter a stop codon in the intronic region that forms the 3’ end of the mRNA. A low proportion of APA events (1.53%) were detected in exonic regions of the ORF, which would lead to nonstop decay due to the lack of a stop codon (reviewed in [28]). In summary, the effect of APA on gene expression is complex, and the global shift of PAS selection that we observed upon LPS stimulation in macrophages is likely to affect the expression of many genes in diverse ways.

When we mimic the effect of LPS on PAS selection by knockdown of *Cpsf5*, secretion of TNF and other pro-inflammatory factors in induced (Fig. 3E-G). This is rather an indirect consequence of altered macrophage polarization, as the mRNAs encoding for these cytokines are not affected by APA (data not shown). In contrast to our observation, CPSF5 was identified as a positive regulator of TNF expression in a CRISPR-Cas9 screen performed in LPS-stimulated RAW264.7 macrophages [29]. Splenic macrophages isolated from myeloid-specific *Cpsf5*-deficient mice also produced less TNF than wild type cells [30]. Moreover, *Cpsf5*-deficient mice were protected from colitis and severe hyperinflammation. Overexpression of *Cpsf5* promoted differentiation of monocytes to macrophages by activation of NF-κB signaling, which led to increased TNF production [31]. While these studies worked with long-term manipulation of CPSF5 expression by knockout or overexpression, we performed short-term depletion of CPSF5 by shRNA-mediated knockdown. The resulting induction of TNF and other pro-inflammatory factors in unstimulated macrophages may activate negative feedback loops that reduce macrophage activity and responsiveness on the long run. This interpretation is also supported by the impaired induction of IL6 and CSF2 upon LPS treatment of CPSF5-depleted cells (Fig. 3G). Therefore, our study revealed an ambiguous role of CPSF5 for macrophage activity.

In RNA-seq data of monocytes isolated from critically ill patients or during sepsis also showed a significant downregulation of *CPSF5* mRNA compared to healthy donors (Fig. 4A). Usage of distal PASs was more heterogeneous in critically ill patients and during early sepsis, with a tendency towards transcript shortening (Fig. 4B). In addition, distal PAS usage shows a negative correlation with *TNF* mRNA expression (Fig. 4C), which is in line with our observation that global transcript shortening can induce TNF secretion (Fig. 3E-G). Although *CPSF5* expression is still reduced during late sepsis, distal PAS usage is comparable to the control condition. This indicates that other factors controlling distal versus proximal PAS usage counteract the repression of *CPSF5*, or that CPSF5 protein levels are restored by post-transcriptional or post-translational mechanisms. In summary, this analysis confirms that PAS selection is affected in monocytes of inflammatory conditions *in vivo*, although with a high variability between individuals, and indicates that compensatory mechanisms restore distal PAS usage at later stages.

## Declarations

### Author contributions

GS and JS designed the study. ML and DL performed experiments. ML and JS analyzed the data. JS, ML and GS wrote the manuscript.

## Acknowledgements

This work was supported by the grant GRK 2727 from the DFG to JS and GS. We would like to thank the NGS Core Facility at the Institute of Clinical Chemistry of the Medical Faculty Mannheim, Heidelberg University, for support with Illumina sequencing

